# LACK OF INTERACTIONS BETWEEN PRENATAL IMMUNE ACTIVATION AND Δ^9^-TETRAHYDROCANNABINOL EXPOSURE DURING ADOLESCENCE IN BEHAVIOURS RELEVANT TO SYMPTOM DIMENSIONS OF SCHIZOPHRENIA IN RATS

**DOI:** 10.1101/2023.01.20.524884

**Authors:** Mario Moreno-Fernández, Marcos Ucha, Raquel Reis-de-Paiva, Alberto Marcos, Emilio Ambrosio, Alejandro Higuera-Matas

**Affiliations:** Department of Psychobiology. School of Psychology. National University for Distance Learning (UNED). Madrid, Spain; UNED International Graduate School (EIDUNED). Madrid, Spain

**Keywords:** Maternal immune activation, THC, adolescence, schizophrenia, animal model

## Abstract

The causality in the association between cannabis use and the risk of developing schizophrenia has been the subject of intense debate in the last years. The development of animal models recapitulating several aspects of the disease is crucial for shedding light on this issue. Maternal infections are a known risk for schizophrenia. Here, we used the maternal immune activation (MIA) model combined with THC exposure during adolescence to examine several behaviours in rats (working memory in the Y maze, sociability in the three-chamber test, sucrose preference as a measure, prepulse inhibition and formation of incidental associations) that are similar to the different symptom clusters of the disease. To this end, we administered LPS to pregnant dams and when the offspring reached adolescence, we exposed them to a mild dose of THC to examine their behaviour in adulthood. We also studied several parameters in the dams, including locomotor activity in the open field, elevated plus maze performance and their response to LPS, that could predict symptom severity of the offspring, but found no evidence of any predictive value of these variables. In the adult offspring, MIA was associated with impaired working memory and sensorimotor gating, but surprisingly, it increased sociability, social novelty and sucrose preference. THC, on its own, impaired sociability and social memory, but there were no interactions between MIA and THC exposure. These results suggest that, in this model, THC during adolescence does not trigger or aggravate symptoms related to schizophrenia in rats.

## Introduction

The study of the relationship between cannabis and schizophrenia has been a recurrent topic in the literature for many years, suggesting that prior cannabis use may be associated with increased risk of psychosis. (Andréasson et al., 1987; De Felice and Laviolette, 2021; Henquet et al., 2005; Ortiz-Medina et al., 2018; Patel et al., 2020; Radhakrishnan et al., 2014). An important developmental perspective in these relationships needs to be considered. Indeed, among cannabis users, the frequent use of the drug and, more importantly, the early onset of use are associated with an elevated risk of transition to psychosis; however, this is not the case for lifetime use (Ehrenreich et al., 1999; Valmaggia et al., 2014). Nevertheless, even if the relationship between cannabis and schizophrenia fulfils several of the standard criteria for causality and the administration of THC to healthy individuals induces schizophrenia-like symptoms such as paranoia, illusions, emotional withdrawal or psychomotor retardation (D’Souza et al., 2004), there are still missing links. To help to shed light on this issue, animal models are essential.

Schizophrenia, per se, is impossible to reproduce in animals, but several models have been developed to mimic the different symptom dimensions of the disease. Some of these models are based on the simulation of maternal infections, an environmental factor associated with an increased risk of schizophrenia diagnosis (Brown and Meyer, 2018), either through exposure to the infective agent itself or through the resulting maternal immune activation (MIA) (Boksa, 2010; Estes and McAllister, 2016). One of the most typical immunogens used in these models is lipopolysaccharide (LPS), a component of the outer membrane of gram-negative bacteria. Indeed, LPS administered during pregnancy in rodents can induce schizophrenia-like symptoms in the adult offspring (Meyer, 2014; Santos-Toscano et al., 2020, 2016; Wischhof et al., 2015). Even if these models are helpful, it has been argued that their validity may be increased by adding a second hit, which would act as a triggering event. Some of these proposed second hits are stressful experiences during adolescence and exposure to cannabinoids during this developmental period (Cattane et al., 2022; Meyer, 2019).

The first animal model combining MIA and cannabinoids was established by Dalton at the beginning of the past decade, using poly I:C to induce MIA in pregnant rats and administering a synthetic cannabinoid (HU210) during adolescence. They found alterations in the integrity of the serotoninergic system in the hippocampus induced by MIA, an effect that they found enhanced by the combination of both factors. (Dalton et al., 2012). Some years later, Hollins and co-workers performed additional experiments with the same hits (poly I:C and HU210), observing, in the left entorhinal cortex, changes in the expression of genes related to schizophrenia, neurotransmission and cellular signalling expression, being the effects more robust in animals exposed to both hits than in those that received only a single hit (Hollins et al., 2016). The relationship between both factors is complex and bidirectional. For example, MIA (induced by poly I:C) alters dopamine transmission but THC during adolescence attenuates some of these effects in Sprague-Dawley rats (Lecca et al., 2019a). The behavioural aspects of these interactions were first tested in a recent work published by Stollenwerk, where the authors used poly I:C to trigger MIA and administered THC through the diet during adolescence and found manifestations of hyperdopaminergic states (measured by amphetamine-induced locomotion) induced by MIA but not caused or enhanced by THC. They obtained similar results with prepulse inhibition and cognitive tests such as the Morris water maze. One of the hits was enough to produce behavioural alterations but without evidence of any synergistic effects (Stollenwerk and Hillard, 2021a). All together, these results again highlight the complex and heterogeneous interaction between MIA and cannabinoid exposure.

To the best of our knowledge, the combination of MIA induced by LPS and an administration of THC during adolescence has never been tested as a model of schizophrenia. Moreover, in the study by Stollenwerk and Hillard, the data were focused on cognitive and emotional processes, but they did not analyse other behavioural dimensions relevant to schizophrenia. Therefore, in this study, we set out to explore different behavioural dimensions that could map to the disparate symptom domains observed in the disease, taking into consideration potential sex-specific effects. Thus, for the assessment of positive symptoms, we used a reality test intended to search for alterations in the accuracy of mental representations of reality (Busquets-Garcia et al., 2017). We used the sucrose preference test and the three-chamber sociability test in order to find evidence of anhedonia or social withdrawal respectively, in an effort to capture some of the negative symptoms of the disease (Ang et al., 2021). For the cognitive symptoms, we used the Y maze test to assess potential impairments in working memory (Hill et al., 2014) and also the widely used prepulse inhibition test (PPI) to detect impairments in sensorimotor gating, typically observed in schizophrenia patients and other animal models of the disease (Khan and Powell, 2018). We also measured somatic growth during adolescence (indexed by the increment in body weight) to obtain a general measure of bodily development.

In addition to study the interactions between both hits in these behavioural dimensions, we also wanted to explore if there were traits in the dams that could predict the appearance or strength of the symptoms in the offspring. There is a great interest to identify such traits. For example, the presence or absence of maternal weight loss after a poly I:C injection has been shown to predict MK-801- and amphetamine-stimulated locomotor abnormalities in the offspring (Bronson et al., 2011a). The results of another study showed that pregnant rats that lost weight following MIA showed increased levels of TNF-α and the most severe behavioural outcome in their offspring compared to controls (S. Missault et al., 2014). However, there are data which point to the fact that intermediate level of baseline immune response to poly(I:C) and an intermediate dose the immunogen are actually most detrimental to the offspring, with higher immune response and poly(I:C) doses conferring resilience to the measured outcomes (Estes et al., 2020a). Clearly, more studies regarding this important topic are warranted. In order to gain a deeper understanding of this issue, in this study we analysed two behavioural traits in the dams (performance in the open field and in the elevated plus maze) and responses to LPS (weight loss and temperature changes after the injection) to compute a susceptibility index that we used to study potential correlations with the behavioural outcome of the offspring.

Contrary to our expectations but in accordance with previous results in the literature, our results do not provide evidence for a triggering effect of adolescent exposure to THC. In addition, we did not find any predictive variables in the dams that could identify future behavioural abnormalities induced by MIA in the offspring.

## Materials and Methods

### Animals

All experimental procedures were previously approved by the Ethics Committee of UNED and the Autonomous Community of Madrid (PROEX 177.4/20). The animals were maintained and handled according to the European Union Laboratory Animal Care Rules (EU Directive 2010/63/EU) and the “Principles of laboratory animal care” were strictly followed.

The study was performed with 30 Sprague-Dawley OFA (Charles-River) rats (15 females and 15 males) that were mated at our centre approximately 3 weeks after their arrival (one male x one female). All animals were maintained at a constant temperature (23 °C) and in a reverse 12h light cycle (lights on at 20:00h) with *ad libitum* access to food (commercial diet for rodents A04; Panlab, Barcelona, Spain) and tap water if not stated otherwise. All experimental procedures were carried between 9:00 and 19:00 and animals were habituated to the test room for 5 minutes 3 days before the first test. Behavioural tests were performed with an illumination of 20-50 luxes.

### Breeding and LPS treatment

Figure 1A shows a diagram depicting the experimental timeline of the behavioural screening performed in the dams and MIA induction.

**Figure 1.**
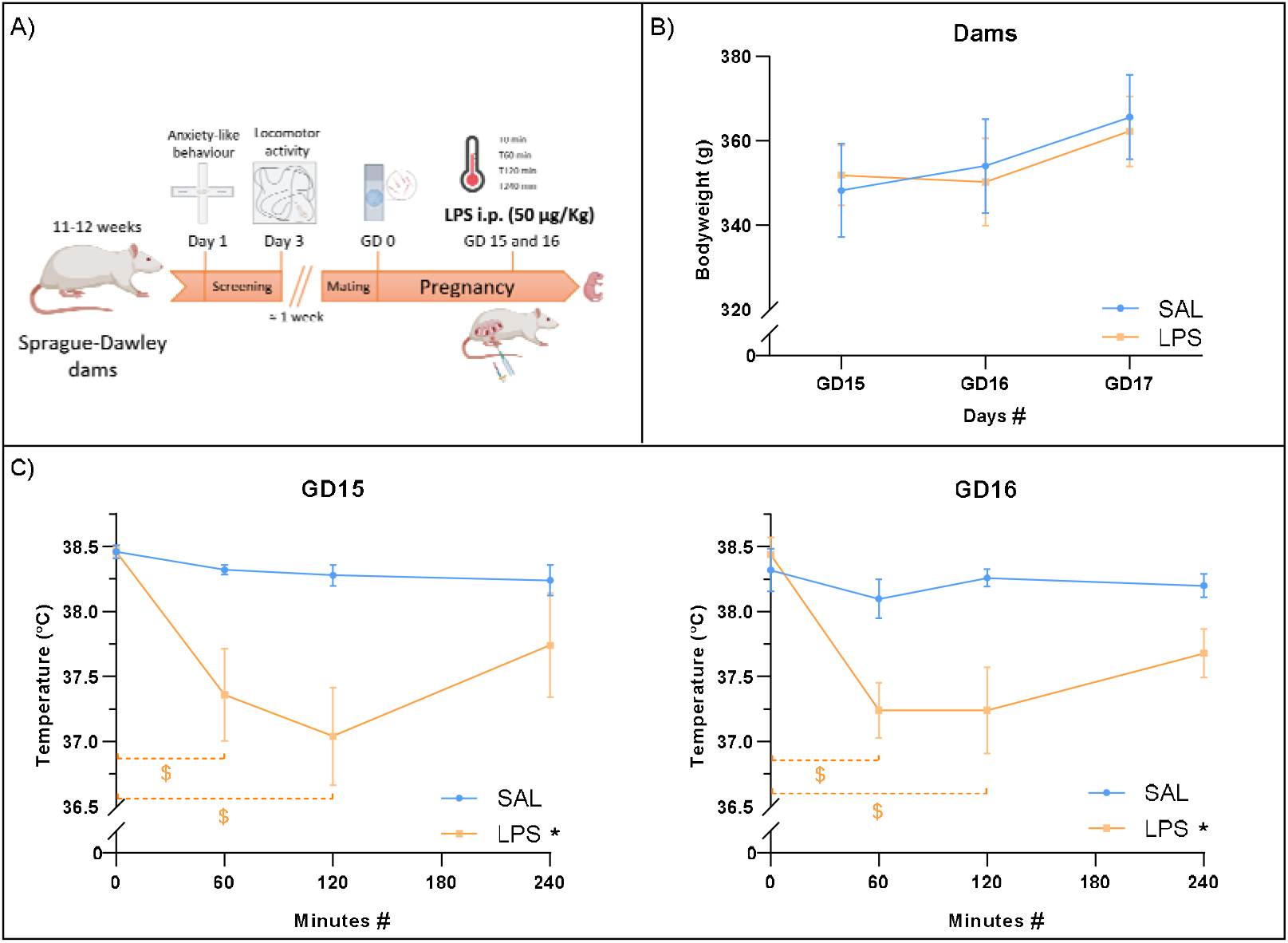
A) Experimental timeline corresponding to behavioural screening of the dams and MIA induction. Elevated plus-maze and open field maze were performed approximately 1 week before mating allowing 1 day between each test. B) Body weight of the dams on the days of injections and one day after is represented as the mean ± SEM. Main intra-subject effect of the days is represented with “#” and intra-subject simple differences between two measures are represented by “$” of the corresponding colour. C) Temperature after LPS injections at gestational days 15 and 16 represented as the mean ± SEM. Main between-subject effect is represented with “*” (p<0.05) and intra-subject effect is represented with “#” (p<0.05). Intra-subject differences of each treatment are represented by “$” of the corresponding colour (p<0.05).

Rats were mated (one male per female per cage) and vaginal smears were taken daily to search for sperm presence as an index to establish gestational day 0 (GD0) when the pregnant rat was isolated. From this moment, body weight was monitored daily. At GD15 and GD16, the pregnant rats were intraperitoneally (ip) injected with 1mL/Kg or LPS 0111:B4 (L2630; Lot# 114M4009V, Sigma-Aldrich) at a dose of 50µg/Kg and volume of 1mL/Kg diluted in sterile saline solution (0.9% NaCl solution; Braun) or saline solution. We reduced the 100µg/Kg dose of our previous studies due to a significant increment of abortions caused after using a new lot of LPS. Rectal temperatures were measured just before the injections and 60, 120 and 240 minutes later as a marker of the MIA. During LPS treatments, 5 pregnant rats reabsorbed the pups.

Between GD21 and GD23, the dams gave birth to 125 pups that were tattooed with Chinese ink in the hind legs to identify prenatal treatment. The litters were then balanced between LPS and saline dams to ensure that each one breeds an equal number of LPS and saline-exposed pups. For the assignment of the pups to their foster mothers, we allowed no more than 1 day of difference between the days of the birth of the rats of each to-be-balanced litter. The resulting cross-balanced litter should not have a difference bigger than 2 pups between males and females.

At postnatal day 22 (PD22) a total of 119 rats were weaned and grouped by sex and treatment in groups of 2-4 rats detailed in Table 1.

**Table 1.** Number of rats per group and treatment <u>THC treatment</u>

| Prenatal | Postnatal | Males | Females |
| --- | --- | --- | --- |
| sal | veh | 15 | 18 |
| sal | THC | 14 | 18 |
| LPS | veh | 13 | 15 |
| LPS | THC | 14 | 12 |
| Total per sex |  | 56 | 63 |
| Total offspring |  | 119 |  |

### THC treatment

Δ^9^-Tetrahydrocannabinol (THC) was obtained from THCPharm (Frankfurt, Germany). The resin was solved first in pure ethanol (30mg/mL) and stored at - 30°C. The working solutions were prepared daily by mixing the stock solution with PEG-35 castor oil (Koliphor; Merck) and saline in a 1:1:18 proportion. All solutions were prepared in siliconized (Sigmacote; Merck) 10 mL vials with a nitrogen-saturated atmosphere and protected from the light. The vehicle aliquots were prepared and treated in the same way, but no THC was added. The THC solution was administered i.p. (3mg/Kg) between PD28 and PD44 every other day (Figure 2A). Body weight was registered every day and 5 days after the end of the administration. The dose and schedule were chosen aiming to mimic the intermittent use of cannabis by human adolescents (Ellgren et al., 2007). The ethanol administered due to THC preparation was approximately 0.0789 g/Kg and does not have significant behavioural effects (Frye and Breese, 1981).

**Figure 2.**
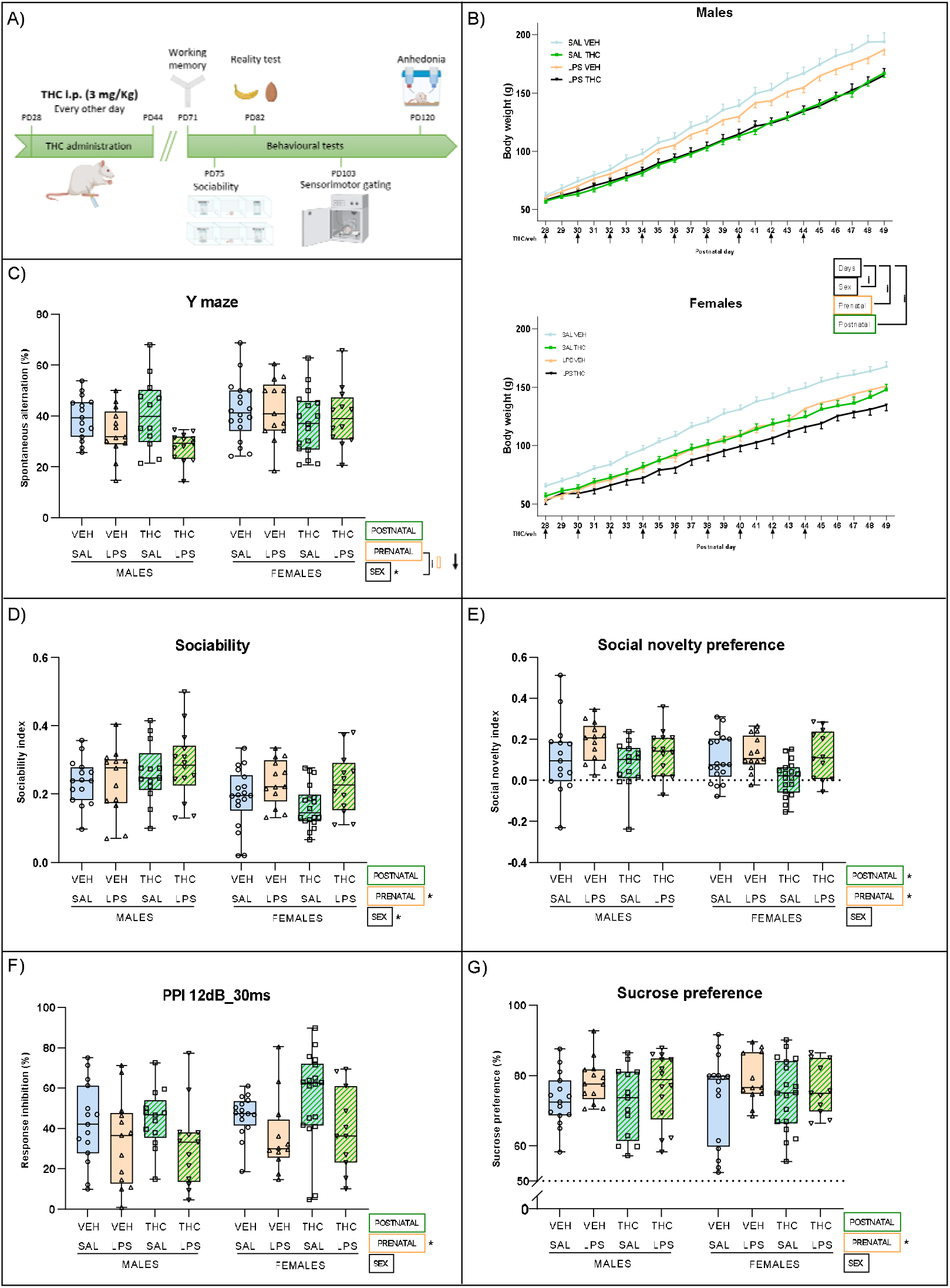
A) Diagram of the THC administration and behavioural tests performed to the offspring. THC (3mg/ Kg) or vehicle i.p. was administered every other day between postnatal day 28 and 44. Body weight was registered through the administration of THC and 5 days after the end. At PD70 behavioural tests were performed in the following order: Y maze, sociability and novelty preference, reality test, prepulse inhibition test and sucrose preference test. B) Body weight change of the offspring through THC administration represented as the mean± SEM. Each THC or vehicle injection is represented with an arrow next to the corresponding postnatal day. C) Spontaneous alteration (percentage of performed triads from total possible triads) assessed with Y maze. D) Sociability index ([interaction t_stranger_ – interaction t_empty_]/ total test time) expressed as percentage evaluated with three-chambered maze. E) Social novelty index ([interaction t_stranger2_ – interaction t_stranger1_]/ total test time) plotted as percentage and analysed with three-chambered maze. F) Prepulse inhibition expressed in percentage and calculated as 1 – [(startle amplitude on prepulse + pulse trial)/ (mean startle amplitude on pulse-alone trials)] x 100. G) Percentage of sucrose preference calculated as [V_sucrose_/ (V_sucrose_+ V_water_)] x 100. Data of all behavioural tests is expressed as mean (bar inside the box) and maximum and minimums values (whiskers).

### Behavioural tests in the progenitor females prior to pregnancy

#### Elevated plus maze

One week before mating, at approximately 11-12 weeks, the anxiety-like behaviour of dams was analysed in the elevated plus maze (Figure 1A). We used an elevated plus-shaped maze made of black methacrylate and expanded PVC (Forex®), elevated 50 cm from the floor and consisting of 2 opposing closed (50×10×50cm) and opened (50×10cm) arms for five minutes, as we previously described (Higuera-Matas et al., 2009) but an arm entry was defined as entry of the centre of the animal. We analysed the data using the following formula: (Open arms time – Closed arms time)/ (Open arms time + Closed arms time) similar to the validated protocol established by Pellow et al., (1985).

#### Locomotor activity

Two days after the elevated plus maze, the locomotor activity in a novel environment was also screened using an open field (50×50×50cm) made of methacrylate. Rats were placed at the centre of the arena and were allowed to freely explore for 40 minutes. The total distance travelled was used as a measure of locomotor activity.

#### Behavioural tests in the offspring

Figure 2A shows a diagram depicting the experimental timeline of the behavioural tests performed in the offspring.

#### Working memory

At PD71 we assessed working memory using a PVC Y-maze, consisting in one longer arm (22×42.5×14cm) and two shorter ones (22×38.5×14cm). The animals were placed on the extreme of the long arm and were allowed to freely explore the entire maze for 10 minutes (Miedel et al., 2017). The total number of entries in each arm and the sequence were recorded. The percentage of alternation was calculated dividing the number of triads with entries into all three arms by the maximum number of possible triads with entries into all three arms (total entries minus two).

#### Social interaction

Social interaction was measured with the three-chamber test at approximately PD75. We used a three chambered maze made of methacrylate consisting in two lateral chambers (39.5×79×39cm) connected to a central one (39.5×79×29.5cm) with guillotine doors. The test consisted of two phases. After 3 minutes of habituation to the central chamber, phase 1 started: wired enclosures (30cm height and 13.5 diameter) were placed on each of the lateral chambers, one was with a same-sex conspecific stranger and the other one was left empty. The guillotine doors were opened, and animals were allowed to explore the entire maze for 8 minutes. Immediately after, phase 2 (social memory/preference test) started: the rat was repositioned within the central chamber with closed doors and a same-sex conspecific unfamiliar rat was placed into the empty wire enclosure. The experimental animal was allowed to explore again the entire maze for 8 minutes. Interaction times, measured as the time that the animal head was inside of a zone surrounding the wired enclosures at a 5cm distance, were recorded in both phases. The social motivation was measured with the Sociability index = (interaction t_stranger_ – interaction t_empty_)/ total test time. Social novelty preference was assessed using the Novelty index = (interaction t_stranger2_ – interaction t_stranger1_)/ total test time.

#### Formation of incidental associations

Around PD82, we analysed the formation of incidental associations with the sensory preconditioning paradigm for impaired reality testing in order to model positive symptoms (such as hallucinations) (Busquets-Garcia et al., 2017). We adapted the protocol described by Koh and colleagues (Koh et al., 2018). During all these procedures, animals had restricted access to water (1h of free access to water in the afternoon and, in the phases which did not involve more time with access to water, 1h in the morning. After three days of habituation to water, the rats were subjected to 4 odour-taste pairing sessions (preconditioning phase) (Figure S1A). For 8 days, they were exposed every morning for 15 minutes on alternate days to two odour/taste combinations: 0.01% almond (benzaldehyde; Sigma) with 5mM NaCl (VWR;) and 0.05 banana (isoamyl acetate; Sigma) with 0.316mM HCl (PanReac Applichem) (Figure S1B). The solution containing the flavour was provided in Falcon™ 50 mL tubes adapted with a sipper tube and weighted before and after the exposure. The solution with the odour stimuli was a filter paper strip soaked with the corresponding solution and placed close to the adapted tube. The concentration of odour and flavoured solutions were chosen to be equally preferred (Busquets-Garcia et al., 2017; Tordoff et al., 2008). The order of presentation of the odour/tasted combinations was counterbalanced between animals (Figure S1C).

The conditioning phase took place the following day. All the experimental groups were exposed for 15 minutes to one of the odours (the odour available during the first preconditioning session) and 5 minutes later they were injected with LiCl (ip, 0.3M in 0.9% NaCl; Sigma). The following day, this process was repeated with the other odour, but only saline solution was administered. During these sessions, the drinking tube was available and filled with plain water. After one recovery day (access to water for 1h in the morning and in the afternoon), animals were exposed for 30 minutes to the solution with the mCS+ (mediated conditioned stimulus): the flavour associated with the LiCl-conditioned odour, and we measured the volume drank by the rats. The next day, we exposed the rats to the mCS-(mediated non-conditioned stimulus): the flavour associated with the other odour and we measured the volume of fluid drank by the rats. The following two sessions were aimed at assessing the direct aversive conditioning, firstly testing the LiCl-conditioned odour (dCS+) by measuring tap water drank in the presence of the odour stimulus, and then testing the other odour (dCS-) one day later.

In order to avoid odour contamination, a room was assigned to each odour and was kept the same during all experiment. The animals remained housed in groups across the experiment and were only single housed the necessary time to be exposed to odours and flavours.

#### Sensorimotor gating

Prepulse inhibition of the acoustic startle response (PPI) was measured at approximately PD103 in a non-restrictive methacrylate cage (28×15×17cm) containing a vibration-sensitive platform (Cibertec), which was calibrated daily calibrated using a 200g weight. The test had a duration of 26 min and was carried out following a sequence previously described by our laboratory (Capellán et al., 2019). Prepulses of 4 or 12 dB above the background noise (65 dB) were used with intervals of 30 or 120 ms between prepulse and pulse (120 dB). Prepulse inhibition was calculated as the %PPI= 1 – [(startle amplitude on prepulse + pulse trial)/ (mean startle amplitude on pulse-alone trials)] x 100. Habituation was assessed by exposing the rats to 6 pulse-alone trials at the beginning and the end of the session (not included in PPI calculations). The percentage of habituation was calculated as [(Mean of first pulse-alone block – Mean of last pulse-alone block)/Mean of first 6 pulse alone block] x 100.

#### Anhedonia

Finally, at about PD120 we carried out a sucrose preference test at PD120 with a protocol based on previous literature (He et al., 2020). The rats had restricted access to water (1h during the afternoon at their home cages), and the experiments tool place during the mornings. The rats had four daily 1 hour sessions in individualized cages with access to two small feeding bottles (one per side) filled with tap water. For the next four sessions, one of the water bottles was replaced by one containing a daily prepared solution of 1% sucrose (Sigma) in tap water. The location of the bottles was counterbalanced between animals and days. Anhedonia was measured using the means of the last 2 days to calculate the percentage of sucrose preference= [V_sucrose_/ (V_sucrose_+ V_water_)] x 100.

#### Video tracking

Animal behaviour was analysed from video tracks with the software ANY-maze Video Tracking System v.6.32 (Stoelting Co.).

#### Data analysis

##### Indexes and correlations

To seek for early markers that may predict some of the deficits seen in this study, we calculated a predictive index for the dams using the data of EPM, locomotor activity, the lowest rectal temperature reached after LPS and weight change after first day of LPS injection. Thus, the index is calculated as follows: DamIndex = [-EPM performance – locomotor activity – Temperature + Weight]. We then calculated an index to group the behavioural tests of the offspring using the average of the Z score of the whole litter for each test and then calculating the severity index as follows Severity Index = [-Ymaze – Sociability – Social novelty preference – PPI – sucrose preference]. Once we obtained both indices we performed several correlation analyses in saline and LPS dams and her litters, using both two indices and their components.

##### Statistical Analysis

IBM SPSS Statistics v.26.0 was used to analyse the data. All results are expressed as the mean ± SEM (standard error of the mean). The interquartile range criterion of SPSS with a p-value<0.05 was applied to identify outliers.

Unless otherwise stated, all experiments were analysed with three-way independent ANOVAs considering “Sex” (male or female), “Prenatal immune activation” (saline or LPS) and “Postnatal THC administration” (vehicle or THC) the between-subject factors. Three-way mixed ANOVAs were used to analyse body weight data taking “Days” with 22 levels as a within-subject factor and the previously mentioned between-subject factors. Three-way mixed ANOVAs were also employed to analyse baseline preferences for the conspecific as compared to the empty compartment in the three chamber test and to verify a the overall preference for the sucrose solution in the sucrose preference test. Furthermore, three-way mixed ANOVAs with two levels (mCS+ and mCS- or dCS+ and dCS-) were used to serach for differences in the mediated and direct tests in the reality test procedure. F-values corrected by Greenhouse-Geisser were used. Significant interactions were followed up by means of simple effects. F-value, effect sizes (partial eta square, η_p_^2^) and degrees of freedom are reported whenever appropriate.

## Results

### LPS administration induces hypothermia in pregnant dams

Temperature along the four sequential measures was differently affected by treatment at GD15 (Time x Treatment interaction *F*_(2.44, 19.54)_ = 4.54, *p* = 0.019,η^2^_p_ = 0.362) and GD16 (Time x Treatment *F*_(2.07, 16.59)_ = 5.01, *p* = 0.018, η^2^_p_ = 0.389) (Figure 2C). Departure from sphericity was at least □=0.51 estimated by Greenhouse-Geisser. In a within-subject analysis we observed a significant decrease of temperature through time caused by LPS on GD15 (*F*_(2.47, 9.86)_ = 6.37, *p* = 0.014, η_p_^2^ = 0.614) and GD16 (*F*_(1.68, 6.72)_ = 8.61, *p* = 0.016, η_p_^2^ = 0.683). Analysing intra-subject differences of the LPS group at GD15 with Tukey-B pos-hoc test adjusted by Bonferroni, we found differences when comparing 0 minutes versus 60 (*p* = 0.013) and 120 (*p* = 0.006). This analysis unveiled the same differences in LPS group at GD16 when comparing 0 minutes versus 60 (*p* = 0.003) and 120 (*p* = 0.022). In addition, the between subjects analysis showed a significant effect of LPS in both days (GD15: *F*_(1, 8)_ = 7.06, *p* = 0.029, η^2^_p_ = 0.469; GD16: *F*_(1, 8)_ = 10.84, *p* = 0.011, η^2^_p_ = 0.575). This hypothermia was not found in saline treated groups (GD15: *F*_(1.54, 6.16)_ = 1.41, *p* = 0.30; GD16: *F*_(1.16, 0.17)_ = 0.683, *p* = 0.47). With the same aim, we analysed body weight changes on GD15, 16 and 17 and found a main effect of the within-subject factor Days (*F*_(1.79, 14.33)_ = 18.26, *p* = 0.000, η^2^_p_ = 0.695) (Figure 2B). From the visual inspection of the data, it would seem that LPS-exposed dams are not gaining weight at the same rate as their saline-controls, but the interaction between days and treatment was not significant (*F*_(1.79, 14.33)_ = 1.41, *p* = 0.274, η^2^_p_ = 0.150).

### THC and MIA independently decrease body weight gain in the offspring

The analysis of the somatic growth of the offspring revealed a main effect of the factor Days in the body weight due to the normal growth of the animals (*F*_(2.31, 228.21)_ = 4663.61, *p* = 0.000, η^2^_p_ = 0.979) (Figure 2B), with a Greenhouse-Geisser estimate of the departure from sphericity of c=0.11. In addition, we found interaction of the factor Days with sex (*F*_(2.31, 228.21)_ = 68.64, *p* = 0.000, η^2^_p_ = 0.409), LPS (*F*_(2.31, 228.21)_ = 4.23, *p* = 0.012, η^2^_p_ = 0.041) and THC (*F*_(2.31, 228.21)_ = 51.98, *p* = 0.000, η^2^_p_ = 0.344). However, we could not find the Days x Sex x LPS x THC interaction (*F*_(1, 99)_ = 0.83, *p* = 0.365). Breaking down the interaction between sex and days we found differences of the body weight along the Days factor in males (*F*_(1.62, 89.23)_ = 1701.59, *p* = 0.000, η^2^_p_ = 0.969) and females (*F*_(1.77, 88.55)_ = 1980.63, *p* = 0.000, η^2^_p_ = 0.975). This results, added to a between subject sex differences (*F*_(1, 99)_ = 13.68, *p* = 0.000, η^2^_p_ = 0.121) point to an expected higher increment of the males body weight along the days. Dissecting the Days by LPS interaction, we observed changes in the body weight along days in saline-descendant (*F*_(1.64, 105.13)_ = 1346.41, *p* = 0.000, η^2^_p_ = 0.955) and LPS-descendant offspring (*F*_(1.40, 57.52)_ = 1202.72, *p* = 0.000, η^2^_p_ = 0.967).

Nevertheless, the body weight increment was lower in the LPS exposed rats than in the saline treated group showed by a between subject analysis (*F*_(1, 99)_ = 5.67, *p* = 0.019, η^2^_p_ = 0.054). Finally, the study of the Days x THC interactions showed an increment in the body weight along the days of vehicle (*F*_(1.69, 93.10)_ = 1696.15, *p* = 0.000, η^2^_p_ = 0.969) and THC groups (*F*_(1.64, 82.07)_ = 1532.97, *p* = 0.000, η^2^_p_ = 0.968). Nevertheless, the between subject analysis revealed alower body weight increment of the rats treated with THC (*F*_(1, 99)_ = 32.11, *p* = 0.000, η^2^_p_ = 0.245). Also, there was a significant main effect of THC administration, showing a diminished weight increment in the rats treated with THC (*F*_(1, 99)_ = 36.04, *p* = 0.000, η^2^_p_ = 0.267).

### LPS prenatal exposure affects working memory in the male offspring

The analysis of the working memory of the offspring using the spontaneous alternation data assessed in the Y maze, revealed a significant main effect of the sex (*F*_(1, 105)_ = 5.38, *p* = 0.022, η^2^_p_ = 0.049) and an LPS x sex interaction (*F*_(1, 105)_ = 6.68, *p* = 0.011, η^2^_p_ = 0.06) indicating a decrease of this parameter in LPS exposed males (*F*_(1, 109)_ = 9.32, *p* = 0.003) (Figure 2C). However, no main effect of LPS (*F*_(1, 105)_ = 3.49, *p* = 0.064) or THC (*F*_(1, 105)_ = 1.98, *p* = 0.162) and no LPS x THC interaction (*F*_(1, 105)_ = 0.44, *p* = 0.509) were found.

### Maternal immune activation increases social interactions in adulthood

In order to study different aspects of social behaviour, we analysed social interactions and social novelty preference. In the first parameter, we evaluate the preference for a conspecific versus an inanimate object in a three-chamber arena. Regarding social interaction, all groups displayed a significant preference for the conspecific (*F*_(1, 108)_=873.78; *p*=0.000 η^2^_p_ = 0.89) (Data not shown). Regarding the sociability index, we observed that females had lower scores in sociability compared to males (*F*_(1, 108)_=10.38; *p*=0.002 η^2^_p_ = 0.088) (Figure 2D). Unexpectedly, we detected a general increment of the sociability index caused in rats exposed to MIA (*F*_(1, 108)_ =4.79; *p*=0.031 η^2^_p_ = 0.042) but, contrary to our expectations, no significant changes were induced by the THC treatment or its interactions with MIA.

Regarding social novelty preference, we found a significant preference of all groups for the novel conspecific (*F*_(1, 105)_ =105.36; *p*=0.000 η^2^_p_ = 0.501) (Data not shown). Interestingly, and in accordance with the results in sociability, a significant increase in social novelty preference was observed in rats subjected to MIA (*F*_(1, 108)_ =8.99; *p*=0.003 η^2^_p_ = 0.077) (Figure 2E). THC decreased this social behaviour component (*F*_(1,108)_ =4.8; *p*=0.031 η^2^_p_ = 0.043) especially in the females, without interacting with MIA experience (*F*_(1, 108)_ =0.622; *p*=0.432).

### Formation of incidental associations: reality monitoring

The three-way ANOVA performed found a main effect of mCS (*F*_(1,49)_ =6.861;*p*=0.012 η^2^_p_ = 0.123) suggesting a decrease in the volume consumed of the mCS solution, however, there were no effects of MIA, exposure to THC during adolescence or their interaction (Figure 3A). When we examined the direct conditioning, the effects of LiCl had already worn off (*F*_(1,44)_ =0.632; *p*=0.431) (Figure 3B), probably due to the amount of time elapsed between conditioning and the test. When including the conditioned odour as a factor (almond or banana) we found a mCS x odour x Prenatal significant interaction (*F*_(1,41)_ =6.645; *p*=0.014 η^2^_p_ = 0.139). More specifically, after performing simple effects analyses, we observed a mCS x Prenatal interaction when almond was the conditioned odour (*F*_(1,28)_ =9.393; *p*=0.012 η^2^_p_ = 0.204). The follow up analysis of this interaction with pairwise comparisons indicated significant differences between mCSs only in saline animals (*F*_(1,28)_ =11.245; *p*=0.002 η^2^_p_ = 0.287) but not in LPS rats(*F*_(1,28)_ =0.187; *p*=0.669). In the case of banana odour, we found a main effect of mCS (*F*_(1,25)_ =8.892; *p*=0.06 η^2^_p_ = 0.262).

**Figure 3.**
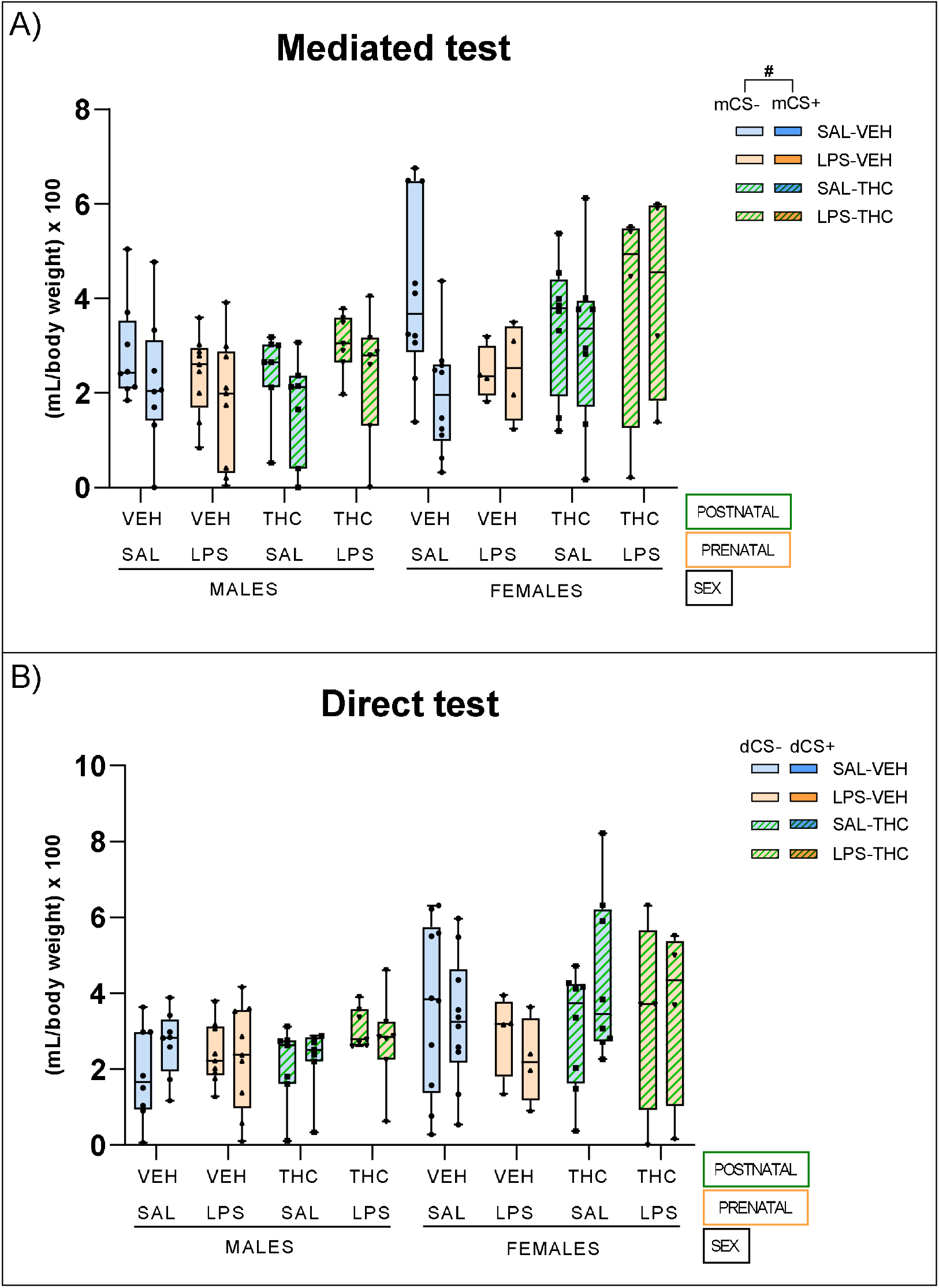
A) Intake volume at the test day of the taste associated to the odour which was conditionated with LiCl (mCS+) or non-conditionated (mCS-). B) Intake volume of at the test day of water in presence of the odour conditionated with LiCl (dCS+) or non-conditionated (dCS-). Data is expressed as mean (bar inside the box) and maximum and minimums values (whiskers). Main intra-subject effect is represented as “#” (p<0.05).

### MIA disrupts sensorimotor gating as assessed by PPI

We observed a significant decrease in the startle response in the 12dB_30ms caused by LPS in both sexes (*F*_(1, 103)_ =10.97; *p*=0.001 η^2^_p_ = 0.096) (Figure 2F) but, contrary to our expectations, we have not detected any interaction with THC treatment (*F*_(1, 103)_ =0.45; *p*=0.506). Similarly, in the 12dB_120ms condition we found a significant effect of LPS in both sexes causing a reduction in the startle response (*F*_(1, 99)_ =5.15; *p*=0.025 η^2^_p_ = 0.049). No more significant results were detected in the rest of PPI conditions (Table 2).

**Table 2.** P and F-values of the different conditions of the prepulse inhibition test. Each condition is represented by the number of dBs above the background noise and the interval between the prepulse and pulse in ms.

| PPI condition | Effect | F-value (degrees freedom) | P-value |
| --- | --- | --- | --- |
| 12dB_120ms | Sex | $F_{(1, 99)} = 0.003$ | 0.954 |
| | MIA | $F_{(1, 99)} = 5.153$ | **0.025** |
| | THC | $F_{(1, 99)} = 0.706$ | 0.403 |
| | Sex* MIA | $F_{(1, 99)} = 1.848$ | 0.177 |
| | Sex*THC | $F_{(1, 99)} = 0.071$ | 0.790 |
| | MIA *THC | $F_{(1, 99)} = 1.666$ | 0.200 |
| | Sex* MIA *THC | $F_{(1, 99)} = 2.363$ | 0.127 |
| 4dB_120ms | Sex | $F_{(1, 86)} = 1.752$ | 0.189 |
| | MIA | $F_{(1, 86)} = 2.136$ | 0.148 |
| | THC | $F_{(1, 86)} = 2.531$ | 0.115 |
| | Sex* MIA | $F_{(1, 86)} = 2.392$ | 0.126 |
| | Sex*THC | $F_{(1, 86)} = 0.299$ | 0.586 |
| | MIA *THC | $F_{(1, 86)} = 0.452$ | 0.503 |
| | Sex* MIA *THC | $F_{(1, 86)} = 0.475$ | 0.492 |
| 12dB_30ms | Sex | $F_{(1, 103)} = 3.047$ | 0.084 |
| | MIA | $F_{(1, 103)} = 10.973$ | **0.001** |
| | THC | $F_{(1, 103)} = 0.720$ | 0.398 |
| | Sex* MIA | $F_{(1, 103)} = 0.052$ | 0.820 |
| | Sex*THC | $F_{(1, 103)} = 0.692$ | 0.407 |
| | MIA *THC | $F_{(1, 103)} = 0.446$ | 0.506 |
| | Sex* MIA *THC | $F_{(1, 103)} = 0.004$ | 0.950 |
| 4dB_30ms | Sex | $F_{(1, 75)} = 3.445$ | 0.067 |
| | MIA | $F_{(1, 75)} = 0.487$ | 0.487 |
| | THC | $F_{(1, 75)} = 0.910$ | 0.343 |
| | Sex* MIA | $F_{(1, 75)} = 0.246$ | 0.621 |
| | Sex*THC | $F_{(1, 75)} = 0.014$ | 0.905 |
| | MIA *THC | $F_{(1, 75)} = 0.027$ | 0.869 |
| | Sex* MIA *THC | $F_{(1, 75)} = 1.744$ | 0.191 |

### MIA increases sucrose preference irrespectively of the sex

Regarding the sucrose preference test, we observed a significant preference for the sucrose solution in all the experimental groups (*F*_(1,103)_ =385.81; *p*=0.000, η^2^_p_ = 0.789) (Data not shown). No significant differences were found in total water (*F*_(1,101)_ =1.29; *p*=0.26) or sucrose consumption (*F*_(1,102)_ =1.40; *p*=0.239). When the sucrose preference between groups was analysed, an unforeseen effect of the LPS was unveiled, consisting on an increased preference for sucrose in this group (*F*_(1, 104)_ = 5.41, *p* = 0.022, η^2^_p_ = 0.049) (Figure 2G). No significant Prenatal x Sex (*F*_(1,112)_ =0.90; *p*=0.765), Prenatal x Postnatal (*F*_(1,112)_ =0.17; *p*=0.896) or Prenatal x Sex x Postnatal (*F*_(1,112)_ =0.114; *p*=0.736) interactions were found.

*Lack of associations between behavioural screening in the dams and severity of deficits in the offspring*

We did not find any significant correlation between the diagnostic indices of the dams and the severity index in the offspring (Table 3).

**Table 3.**
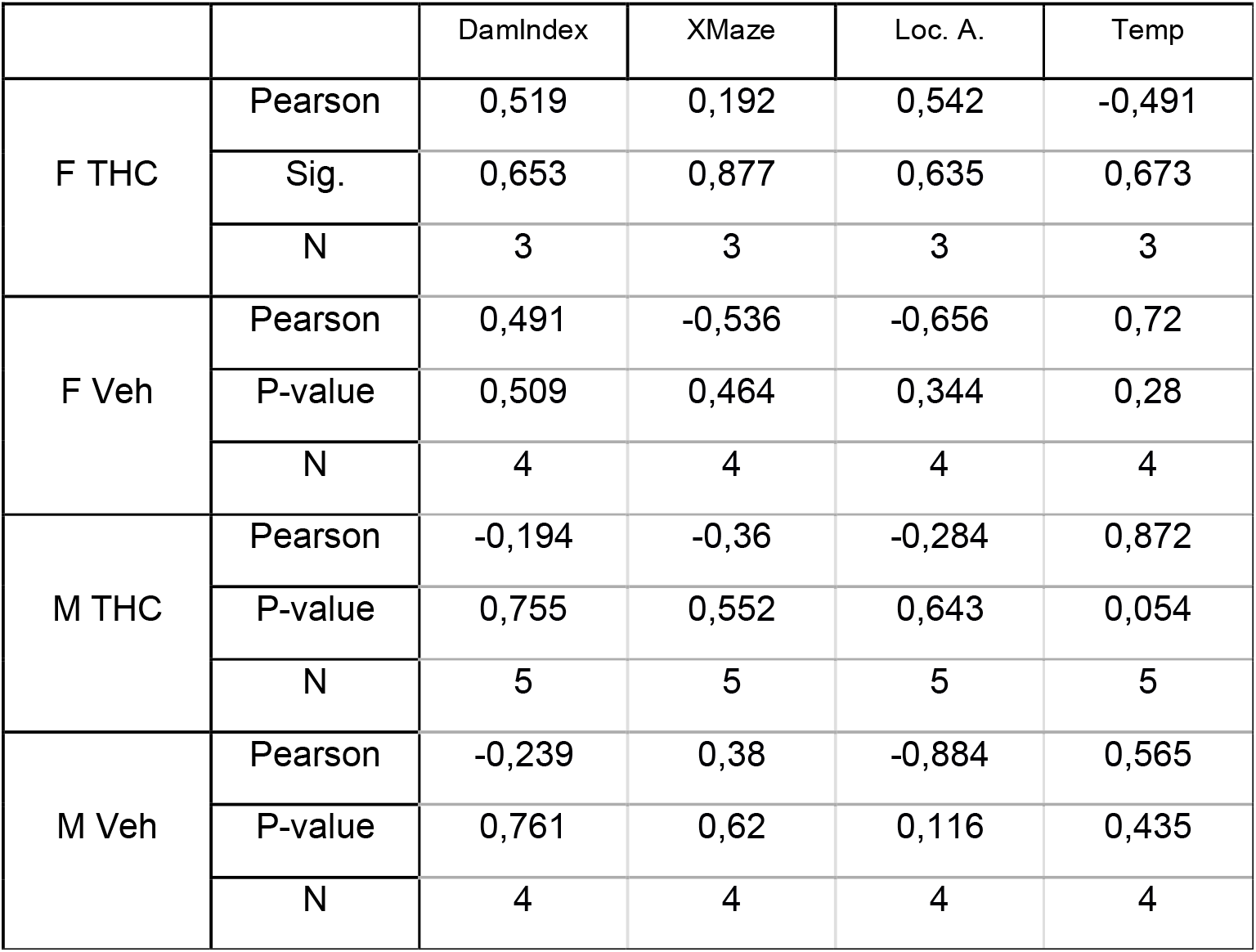
Pearson product-moment correlation tests between the behavioural index of the dams (and its components) and the diagnostic index of the of offspring (according to sex and Vehicle/THC exposure). Loc. A. (Locomotor activity).

## Discussion

The data presented here does not support a two-hit hypothesis of schizophrenia wherein MIA, as a susceptibility factor, would act in synergy with cannabinoid exposure during adolescence (the triggering hit) to generate the different symptoms associated with the disease. Moreover, we could not detect predictive variables which could identify subjects at higher risk of developing the array of behavioural disturbances observed in schizophrenia or related neurodevelopmental disorders.

Although we found no evidence of synergy between MIA and THC exposure, both hits, separately, had interesting effects on somatic growth and adult behaviour.

### Effects on body weight

We found that MIA and THC exposure during adolescence independently, reduced body weight between PD28 and PD50 (Figure 2B); hence, this effect was still present even after 6 days of THC washout. The data available regarding the effects of MIA induced by LPS on body weight regulation in the offspring are inconclusive. While some studies have evidenced a reduction in body weight (Bakos et al., 2004; Izvolskaia et al., 2016), others found no evidence of disruption (Harvey and Boksa, 2014) and some other works report an increment in body weight (Bakos et al., 2004; Ni et al., 2022; Wei et al., 2007). The reasons for these discrepancies still need to be clarified. For example, the study by Harvey and Boksa showed no evidence of altered body weight following MIA in the offspring, using the same rat strain, dose, way of administration and gestational days as us. Moreover, these authors measured body weight at PD30 and PD60, which covers the developmental period we examined. Therefore, the rat strain, dose, way of administration, the developmental period for LPS administration and measurement time points are not explicative factors.

We also found that THC administration decreased body weight, an effect that persisted after treatment cessation. THC exposure during adolescence is known to reduce body weight in animal models (Keeley et al., 2015; Klein et al., 2011; Rubino et al., 2008) (although these data do not seem to apply to humans, see (Jin et al., 2017) for an example)) so the results presented here are in agreement with previous reports. Noteworthy, we have not detected significant interactions of MIA or THC administration with the sex of the animals or between them. However, MIA effects on body weight seem to be more robust in females.

### Cognitive symptoms: working memory and sensorimotor gating

MIA was associated with deficits in these two cognitive domains. Working memory, as assessed in the Y maze, was impaired in the offspring exposed to MIA, which is in agreement with similar alterations documented before (da Rosa et al., 2022; Wischhof et al., 2015). Interestingly, this effect was restricted to the male offspring, as previously reported by our laboratory, when we used a T maze to explore working memory (Santos-Toscano et al., 2016). It is tempting to invoke a role for oestrogens, or the oestrogen receptor, in particular, to explain these sex-related differences (Arnold and Saijo, 2021). Still, other explanations, such as differences in the oxidative stress response (Tower et al., 2020) or antioxidant defence systems (Julicher et al., 1984), should not be disregarded.

THC did not affect working memory, and there was no evidence of synergy between both hits. Other studies have shown no alterations in Y maze performance following adolescent THC (Cadoni et al., 2015), and THC self-administration during adolescence even slightly improved working memory in a delayed-match-to-sample task. In general, the effects of cannabinoid exposure during adolescence on cognition are complex and critically depend on the type of administration (passive or volitional), age of exposure, memory task used, or the cannabinoid employed (natural or synthetic) (Zamberletti and Rubino, 2022).

Regarding PPI, sensorimotor gating has traditionally been used as a proxy to evaluate positive symptoms in human patients and animal models of schizophrenia-like behaviours (Schmidt-Hansen and Le Pelley, 2012; Wynn et al., 2004). Our results showed PPI reduction in both male and female MIA rats (Figure 2F), in accordance with previous studies (Wischhof et al., 2015); however, in agreement with previous literature, we found no detrimental effects of THC exposure on PPI. Importantly, we found no interaction between MIA and cannabinoid exposure during adolescence, suggesting that there are no summative or synergistic actions between both hits in this model, as reported with other two-hit models (Garcia-Mompo et al., 2020; Guma et al., 2023; Rodríguez et al., 2017; Stollenwerk and Hillard, 2021a).

### Negative symptoms: sociability, social novelty preference and anhedonia

Negative symptoms of schizophrenia are associated with a reduced quality of life and functional status (Möller, 2016). We analysed different social parameters using the three-chamber maze to evaluate features that may mimic some aspects of this domain. Our data show a general increase in social interaction caused by MIA and lower basal social interaction in females compared to males (Figure 2D-E). This higher sociability in males has been reported in the past (Cossio et al., 2020). Previous data on the effects of MIA on social interaction are not conclusive. While several studies found significant social deficits (Lee et al., 2021; Talukdar et al., 2020), other authors could not find differences in this respect (Batinić et al., 2016). Furthermore, cytokines, such as IL-17a, associated with MIA also seem to rescue sociability deficits in genetic mouse models of autism spectrum disorder and MIA offspring (Reed et al., 2020). The different doses and administration times may partially explain the significant variability of the data. In addition, it is known that the different batches of immunogenic agents (LPS or Poly I:C) induce immune responses of variable strength, generating more inconsistency in the data from the different laboratories (Mueller et al., 2019). Finally, the differences in the species of the experimental animals used standard (mice or rats) could contribute even more to the absence of homogeneous data.

Anhedonia has traditionally been a main negative symptom of schizophrenia (DSM-V, 2013; RADO, 1953). We performed a sucrose preference test to seek similar features in our model. Our data shows a general increase in sucrose preference in MIA rats of both sexes (Figure 2G). Contrary to our predictions, this may indicate an enhanced hedonic reaction caused by LPS exposure during pregnancy. This is against the anhedonia caused by Poly I:C in rats observed by Missault (S. Missault et al., 2014). However, in the last decade, the heterogeneity of the anhedonia dimension across the major psychiatric disorders has raised the need for a more complex definition of anhedonia. “The lack of ability to experience pleasure” is being replaced by “impairment in the ability to pursue, experience and/or learn about pleasure” (Thomsen et al., 2015). This has allowed a better understanding of anhedonia in schizophrenia, as recent evidence postulates that the capacity to anticipate pleasure is the actual psychological process impaired in these individuals (Šagud et al., 2019) and not the consummatory dimension of pleasure (Fortunati et al., 2015). Altogether, this evidence demands a more detailed study of anhedonia in all the domains of this dimension in future animal models.

### Positive symptoms: formation of incidental associations, reality monitoring

We have evaluated the accuracy of mental representations of reality, a feature that may mimic or underlie features like hallucinations in schizophrenia (Busquets-Garcia et al., 2017; McDannald and Schoenbaum, 2009). We did not observe differences between groups and only found a general decrease in the preference for the mCS+, suggesting that in this specific protocol in rats, a preconditioning phase consisting of 4 pairings per compound is insufficient for the animals to separate the mental representation of the two stimuli clearly. Previous studies have successfully shown effective reality monitoring in control animals using the same number of pairings during preconditioning that we used (McDannald et al., 2011). Still, the protocol used in these previous experiments was based on operant responses rather than classic conditioning, which may explain the diverging results. Future studies should test a higher number of pairings to explore further if there is an impairment in reality monitoring following MIA, adolescent cannabinoid exposure or their combination.

### A search of behavioural predictors of symptom severity

There is an increasing interest in finding markers of immune response after maternal infection that could predict if schizophrenia or related neurodevelopmental disorders will ensue in the offspring. MIA models have been crucial in this endeavour and have focused on the baseline immune reactivity to the immunogen, weight loss or rectal temperature changes after the immune challenge (Bronson et al., 2011b; Estes et al., 2020b; Lins et al., 2018; S Missault et al., 2014; Vorhees et al., 2012). In an attempt to contribute to this search, we decided to explore if behavioural traits in the rats, measured before pregnancy and immune challenge (open field and elevated plus maze performance) and also the classical measures of immune reactivity following a challenge (weight loss and temperature changes), could be combined to compute a predictive index. Contrary to our expectations, these behavioural markers did not predict symptom severity either in combination, as a compound index, or individually. However, the low sample size (we averaged the score of every rat born per dam to obtain a single measure and correlate it with the corresponding value of their mother, losing statistical power in the process) is likely to explain this lack of results.

### Conclusions

In this work, we have shown that MIA and THC exposure during adolescence do not interact to precipitate schizophrenia-related symptoms in rats, at least not in the behavioural dimensions that we have examined. This adds to the growing number of studies that have experimentally tested the two-hit hypothesis and have found no evidence for synergy between the two hits (Guma et al., 2023; Lecca et al., 2019b; Stollenwerk and Hillard, 2021b). Clearly, we are still at an early stage of research using these two-hit models before we can unmistakably dismiss a precipitating role of cannabis consumption during adolescence in the onset of schizophrenia. Future studies should test higher doses of THC and examine brain function parameters and different brain neurochemistry elements to understand how these two proposed risk factors may interact.

**Figure S1.**
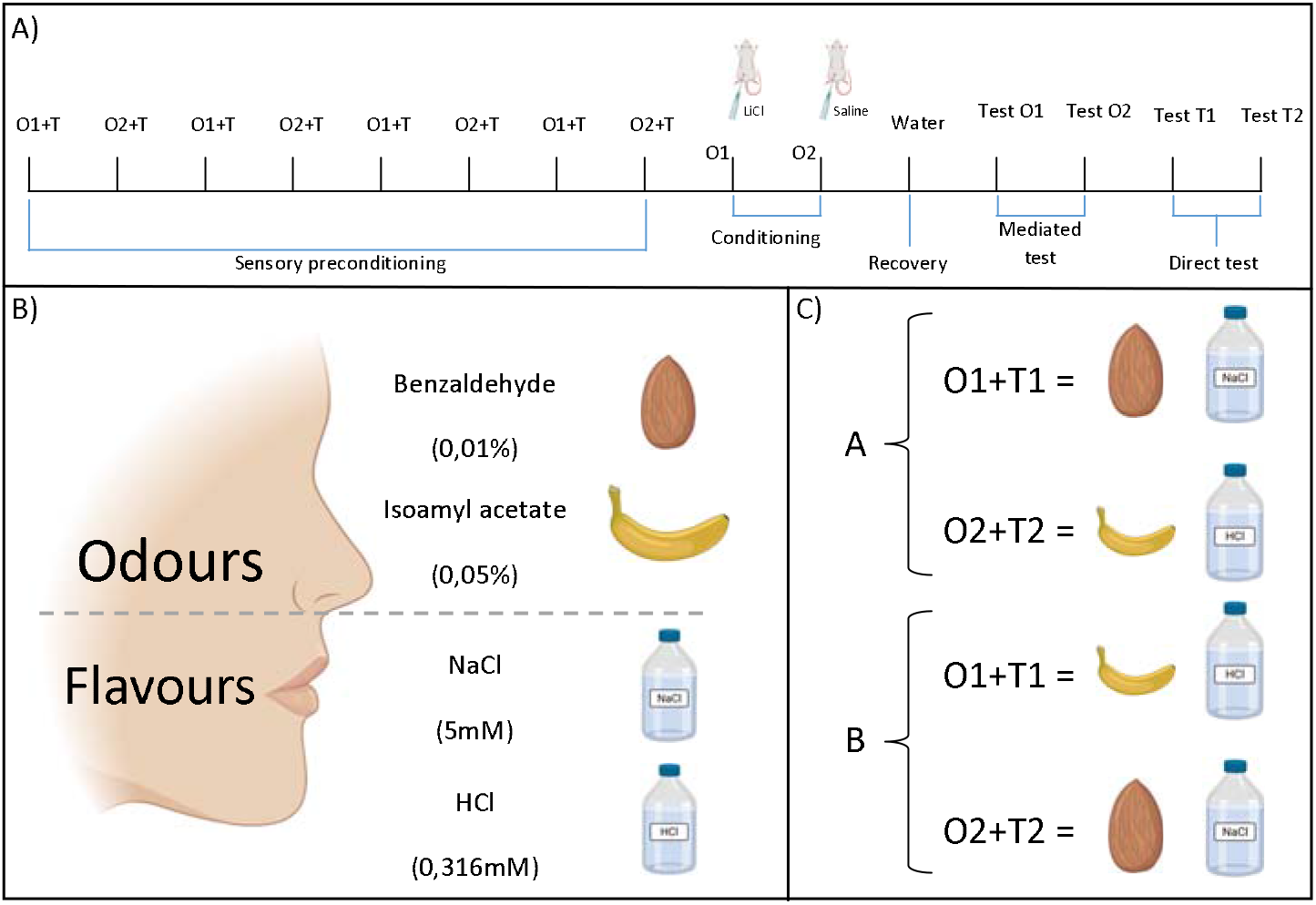
Diagram of reality test. A) Temporal scheme of the protocol. Animals were exposed to O1+T1 and O2+T2 at alternate days to produce a weak association between stimuli. After 4 exposures of each pair, O1 was aversively conditioned with LiCl and O2 with saline as a control. After one day of recovery each taste (mediated conditioned stimuli) and each odour (direct conditioned stimuli) were tested. Each vertical line represents one day and always is aversively conditioned O1 with LiCl. B) Reagents and concentrations used for odours and tastes C) Counterbalancing used to avoid odour preferences. Half of the animals had almond and NaCl as O1 and T1 and the other half had banana and HCl as O1 and T1.

## Acknowledgments

We would like to thank Rosa Ferrado for her excellent technical assistance.

